# Human activity, not environmental factors, drives *Scedosporium* and *Lomentospora* distribution in Taiwan

**DOI:** 10.1101/2024.11.07.622530

**Authors:** Hsin-Mao Wu, Yu-Hsuan Fan, Guan-Jie Phang, Wen-Ting Zeng, Khaled Abdrabo El-Sayid Abdrabo, Yu-Ting Wu, Pei-Lun Sun, Ying-Hong Lin, Yin-Tse Huang

## Abstract

*Scedosporium* and *Lomentospora* species are emerging fungal pathogens capable of causing severe infections in both immunocompetent and immunocompromised individuals. Previous environmental surveys have suggested potential associations between these fungi and various soil chemical parameters, though the relative influence of human activity versus environmental factors has not been systematically evaluated. Here, we conducted a comprehensive survey of 406 soil samples from 132 locations across Taiwan, analyzing fungal abundance alongside soil physicochemical parameters and the Human Footprint Index (HFI). We recovered 236 fungal isolates comprising 10 species, with *S. boydii* (32.2%), *S. apiospermum* (30.9%), and *S. dehoogii* (14.4%) being the most prevalent. The highest fungal burdens were observed in urban environments (up to 1293 CFU/g), particularly in public spaces and healthcare facilities. Statistical analysis revealed a significant positive correlation between fungal abundance and HFI (r = 0.125, p = 0.013), while soil chemical parameters including nitrogen, carbon, pH, electrical conductivity, and various base cations showed no significant associations despite their wide ranges. These findings indicate that anthropogenic disturbance of environments, rather than soil chemistry, is the primary driver of *Scedosporium* and *Lomentospora* distribution in Taiwan. This understanding holds important implications for predicting infection risks and developing targeted public health strategies, particularly in rapidly urbanizing regions. Future studies incorporating more specific indicators of human impact may further elucidate the mechanisms underlying these distribution patterns.

**Lay Abstract:** We found that human activities, rather than soil properties, determine where *Scedosporium*/*Lomentospora* fungi distribute in Taiwan. These fungi are more abundant in urban areas than less-human-disturbed environments, suggesting increased infection risks in densely populated regions.

## Introduction

*Scedosporium* and *Lomentospora* species are clinically significant fungal pathogens capable of causing a wide spectrum of human infections. These range from localized infections like mycetoma following traumatic inoculation in immunocompetent hosts, to severe disseminated infections in immunocompromised patients, particularly those with hematologic malignancies or transplant recipients.^1,2^ In cystic fibrosis patients, these fungi rank second among filamentous fungi colonizing the airways,^3^ and they are particularly concerning due to their ability to cause allergic bronchopulmonary disease and potential progression to invasive infection following lung transplantation.^4^ The mortality rates are notably high, reaching up to 80% in cases of disseminated infection and 75% in neutropenic patients with invasive disease.^2^ Due to their intrinsic resistance to most available antifungal drugs and increasing clinical significance,^5–7^ *Scedosporium*/*Lomentospora* species have been recently included in the World Health Organization’s first-ever Fungal Priority Pathogens List (FPPL),^8^ highlighting their global importance as emerging pathogens requiring urgent attention and research focus.

Environmental surveys conducted worldwide have provided important insights into the ecological preferences of these fungi. Studies in Austria and the Netherlands found no *Scedosporium* species in natural habitats such as forests and sand dunes, but reported high recovery rates from industrial areas (up to 641 CFU/g soil), parks and playgrounds (388 CFU/g soil), and agricultural lands (25 CFU/g soil).^9^ In France, the highest densities were found in human-impacted areas, with *S. dehoogii* predominantly recovered from industrial areas and *S. aurantiacum* from agricultural regions.^10^ Australian surveys revealed *S. aurantiacum* (54.6% of isolates) and *L. prolificans* (43%) as predominant species in urban environments, with significantly higher fungal burdens in areas within 3 km of city centers compared to rural sites.^10^ In Thailand, environmental surveys in human-impacted public parks found high abundance and diversity of Scedosporium species, with S. apiospermum as the predominant species, accounting for 86.6% of isolates in Bangkok parks. Broader surveys across regions with dense human activity and tourism further confirmed S. apiospermum’s dominance, revealing significant genetic diversity among isolates, including potential novel species.^11,12^ Mexican surveys across 25 states found these fungi primarily in areas of high human activity including urban gardens (49%), industrial parks (24%), and sports parks (14%).^13^ Notably, these studies consistently demonstrated a strong association between human activities and the presence of *Scedosporium*/*Lomentospora* species, while also revealing geographical variations in species distribution.

The distribution and abundance of *Scedosporium*/*Lomentospora* species are strongly influenced by environmental factors, with nutrient availability playing a particularly crucial role.^9,14^ These fungi show clear preferences for nitrogen-rich environments, especially those with elevated nitrate levels, helping explain their prevalence in fertilized agricultural lands and areas with intensive animal activity.^9^ Phosphorus availability also appears important, with some studies reporting optimal abundances in phosphorus-rich soils (200-300 ppm assimilable P), particularly in wastewater treatment systems.^14^ Organic matter content has been consistently associated with higher fungal populations across different geographical regions.^14^ Temperature has emerged as another key factor that not only directly affects fungal growth but also modulates the impact of other environmental variables. While temperature alone shows modest effects on *Scedosporium* populations, warmer conditions (around 25°C) significantly enhance the influence of both nutrient availability and environmental pollutants on fungal abundance.^15^ Other environmental factors including pH and electrical conductivity have also been implicated in shaping *Scedosporium* distribution. While optimal pH ranges between 6.1-7.6 have been widely reported,^9,11,14^ populations can thrive across broader pH conditions (4.9-8.5) in some regions.^10^ Electrical conductivity alone has been shown to explain nearly 20% of variance in abundance, suggesting these fungi are well-adapted to moderately saline conditions.^14^ Additionally, hydrocarbon contamination, particularly from diesel fuel, appears to strongly influence population densities, with some studies reporting substantially higher fungal counts in polluted soils.^15–17^ However, these reported relationships between environmental factors and fungal abundance have largely been based on correlation studies from field observations, with Rainer and Eggertsberger providing the first experimental validation for the effects of temperature, nitrate, and diesel contamination on *Scedosporium* populations.^15^ Their controlled studies demonstrated that these factors can act synergistically, with temperature amplifying the effects of both nitrogen availability and hydrocarbon contamination. Nevertheless, the strong associations with human-impacted environments suggest that anthropogenic factors, rather than soil chemistry alone, may be the primary drivers of *Scedosporium*/*Lomentospora* distribution and abundance in natural ecosystems.

In Taiwan, our previous study revealed high recovery rates of *Scedosporium* species from soil samples across urban and hospital areas in Taiwan.^18^ To build upon these findings and better understand the environmental factors governing the distribution of these medically relevant fungi, we conducted an expanded environmental survey incorporating three key components: 1) more comprehensive geographical coverage including eastern Taiwan, 2) systematic analysis of soil physicochemical parameters, and 3) quantitative assessment of human impact using the Human Footprint Index (HFI). This multifaceted approach aims to delineate the relative importance of anthropogenic influence versus edaphic factors in determining the abundance and distribution of *Scedosporium* and *Lomentospora* species across Taiwan’s diverse landscapes.

## Materials and Methods

### Sample sites

A total of 132 distinct locations across Taiwan were investigated in this study, comprising the 79 previously sampled locations in Huang et al. (2023)^18^ and an additional 53 new sampling sites, particularly strengthening the coverage in eastern Taiwan (Fig. 1, Supplementary Table 1). Land use type classification for all sampling sites was based on the Land Use Investigation (LUI) Map from The National Land Surveying and Mapping Center (NLSC) (https://www.nlsc.gov.tw/), Taiwan. Across the 132 locations, a total of 406 soil samples were collected, resulting from 1-17 replicates per location to account for spatial variability. At each location, 1-4 sampling units were randomly selected within a square meter. For each sampling unit, soil was collected from five points with 10-20 cm intervals from the center using a metal spatula sterilized with 70% ethanol between samples to avoid cross-contamination. Soil was sampled at a depth of 7-10 cm beneath the soil surface, pooled in a sterile plastic bag, and gently mixed, then transferred to a 50 ml sterile Falcon tube. All soil samples were stored at 4°C until processing.

**Figure 1.**
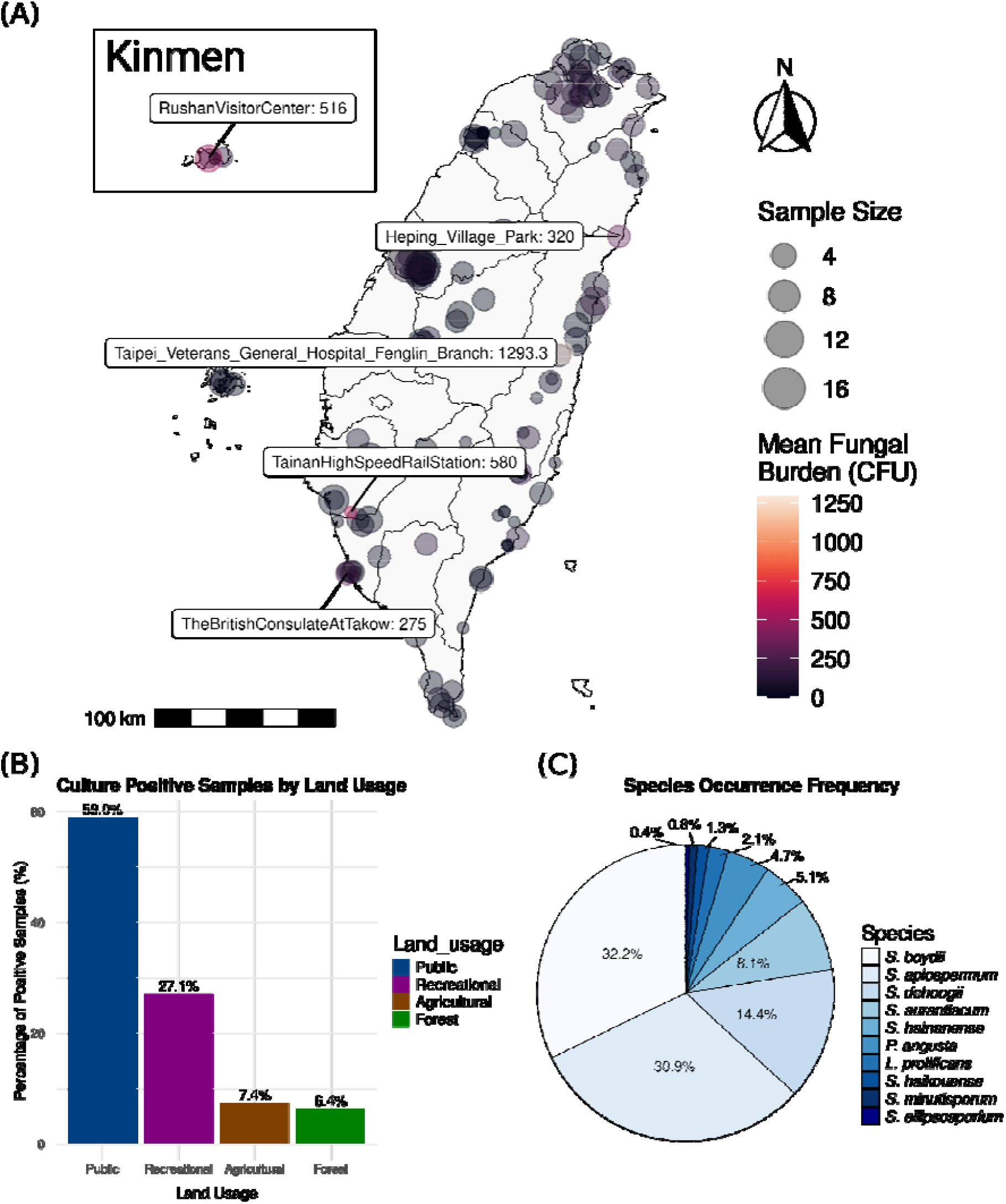
Distribution and prevalence of *Scedosporium*/*Lomentospora* species across Taiwan. (A) Mean CFU of *Scedosporium*/*Lomentospora* species at each sampling location, with locations showing mean CFU > 200 labeled; circle sizes represent sample sizes, and values indicate CFU per gram. (B) Percentage of culture-positive samples across land use types in Taiwan. (C) Species distribution among 236 isolates, displaying occurrence frequencies for each *Scedosporium*/*Lomentospora* species.

### Isolation and Quantification of Fungal Burden in Soils

For each soil sampling unit, 10 g of soil was air dried at room temperature for 1-2 days, placed in a 100 ml sterile Erlenmeyer flask and homogenized with 90 ml sterilized water containing 1-2 drops of Tween 20. The mixture was thoroughly vortexed and left to stand for 15 min. A total of 250 μl of suspension was inoculated onto five plates of Dichloran Glycerol (DG18) Agar Base (Himedia, India) supplemented with chloramphenicol. Plates were incubated at 37°C for 5-7 days and examined at intervals for the emergence of characteristic colonies of *Scedosporium*/*Lomentospora*. All *Scedosporium*/*Lomentospora* colonies on five plates of each sampling unit were counted, from which the fungal burdens were determined based on the mean value of colony forming units (CFU)/g of soil dry weight.

### Molecular Identification of Fungal Isolates

For molecular identification, we amplified the internal transcribed spacer (ITS) region using primer pairs as previously described.^19,20^ PCR amplification was performed with a two-step barcoding approach according to Herbold et al. (2015).^21^ The first-step PCR reaction mixture volume was 25 μl, consisting of 1 μl DNA template, 1 μl of each primer, 12.5 μl of 2X Tools Supergreen PCR master mix, and 9.5 μl of ddH2O. Amplification was carried out with initial denaturation at 95°C for 2 min, 25 cycles with denaturation at 95°C for 30 s, annealing at 64-66°C for 30 s, primer extension at 72°C for 1 min, and final extension at 72°C for 7 min. The second-step PCR used 1 μl of the first-step product as template with pre-mixed barcoded primers. The products were purified using Ampure Xp Beads with 0.4X product to bead volume ratio. Sequencing was carried out on an Oxford Nanopore Technologies GridION sequencer using the manufacturer’s library preparation kit (SQK-LSK110).

We reconstructed the phylogeny of the isolated species using the ITS sequences. Sequences of type specimens of *Scedosporium* and *Lomentospora* species were retrieved from NCBI Genbank. Sequences of *Petriellopsis africana* (CBS 311.72), *Petriella setifera* (CBS 385.87), and *P. sordida* (CBS 144612) were selected as the outgroup. Sequences were aligned using MAFFT.^22^ Phylogenetic informative regions were selected using ClipKIT^23^ with the gappy strategy. Phylogenetic trees were reconstructed using IQ-TREE2^24^ with recommended partition parameters inferred using ModelTest-NG.^25^ Trees were visualized using Interactive Tree of Life version 4.^26^

### Soil Chemical Properties Analysis

All soil chemical analyses were performed on air-dried soil samples that passed through a 2-mm sieve. Each analysis was performed in duplicate, and mean values were used for subsequent statistical analyses.

Total nitrogen (N) in soil samples was determined using the semimacro Kjeldahl method with a KB-8 digestion block and VAP 300 steam distillation system (Gerhardt GmbH, Germany). Soil organic matter (OM) content was assessed via wet oxidation following the Walkley-Black procedure. Water content was determined by oven-drying soil samples at 105°C until constant weight was achieved. Soil pH and electrical conductivity (EC) were determined using a pH/EC/TDS/Temperature Portable Meter HI9814 (HANNA Instruments,Woonsocket, RI, USA) in a 1:2.5 soil: water suspension.

Available phosphorus was measured using the Bray-1 method with spectrophotometric determination. Exchangeable cations (Na+, K+, Mg2+, and Ca2+) were assessed by atomic absorption spectrophotometry (AAS) using a Z 5300 instrument following the manufacturer’s recommendations (Hitachi—Science & Technology, Tokyo, Japan). For this analysis, soil samples were extracted with 1 M ammonium acetate (pH 7.0), and the extracts were analyzed for cation concentrations.

## Statistical analyses

Statistical analyses were performed using R version 4.1.0 with the ggplot2,^27^ dplyr,^28^ vegan,^29^ ggpubr,^30^ sf, ggrepel,^31^ viridis,^32^ and ggspatial^33^ packages. To assess the distribution and prevalence of *Scedosporium* and *Lomentospora* species across different land usage types (Public, Recreational, Agricultural, and Forest), samples were classified as culture-positive if they contain any detectable colony-forming units (CFUs) of these fungi.

The percentage of positive samples was calculated for each land usage category, and species occurrence frequency was determined by calculating the proportion of each identified fungal species among total detections.

Beta diversity analyses were conducted to quantify differences in species composition between samples using a Bray-Curtis dissimilarity matrix. Variability in species composition among land usage types was examined through permutational analysis of multivariate dispersions (PERMDISP) using the betadisper() function in the vegan package, with land usage as the grouping factor. An analysis of variance (ANOVA) on the betadisper model tested for significant differences in dispersion across land use categories. Post hoc pairwise comparisons were performed using Tukey’s Honest Significant Difference (HSD) test, and results were visualized with boxplots of distances to group centroids.

To map the distribution of fungal burden across Taiwan, mean fungal burden per location was calculated by aggregating CFU values of *Scedosporium* and *Lomentospora* species across samples, along with the total number of samples per site. Geographic visualization was performed using spatial packages, with point size representing the sample count per location and color intensity indicating the mean fungal burden. Labels were added for sites with mean fungal burdens exceeding 200 CFUs to highlight locations with higher fungal loads.

To investigate the relationships between fungal burden, Human Footprint Index (HFI), and various chemical and edaphic factors, CFU data for multiple fungal species were analyzed alongside soil chemistry data. CFU counts were log-transformed to normalize the data, and the average CFU value per sample was calculated to represent the mean fungal burden.

Correlations between mean fungal burden and chemical factors—including the HFI [retrieved from Human Footprint, Last of the Wild Project, Version 3, 2018 Release],^34,35^ total nitrogen, total carbon, organic content, water content, pH, electrical conductivity (EC), available phosphorus, sodium, potassium, magnesium, and calcium—were evaluated using generalized linear models (GLMs) with a Gaussian family. For each factor, correlation coefficients, p-values, and sample sizes were computed to assess the strength and statistical significance of associations.

## Results

### Isolation and Quantification of Fungal Burden in Soils

A total of 236 fungal isolates belonging to 10 different species within the *Scedosporium* and *Lomentospora* species were recovered from soil samples collected across Taiwan (Figure 1). The most frequently isolated species was *S. boydii* (76 isolates, 32.2%), followed by *S. apiospermum* (73 isolates, 30.9%), and *S. dehoogii* (34 isolates, 14.4%). Less commonly isolated species included *S. aurantiacum* (19 isolates, 8%), *S. hainanense* (12 isolates, 5%), *P. angusta* (11 isolates, 4.7%), and *L. prolificans* (5 isolates, 2.1%). *S. haikouense* was rarely isolated (3 isolates, 1.3%), *S. minutisporum* (2 isolates, .0.8%), and *S. ellipsosporium* (1 isolate, 0.4%). The newly generated sequences from this study have been deposited in NCBI GenBank under accession numbers PQ569163 to PQ569295, encompassing 133 isolates. For additional sequence accession numbers, refer to Huang et al. (2023).^18^

Fungal burdens varied considerably among sampling sites and land usage types (Figure 1). The highest fungal burdens were observed in urban environments, particularly in public areas and recreational sites. Notable examples included Taipei Veterans General Hospital Fenglin Branch (up to 1293 CFU/g), Tainan High Speed Rail Station (up to 580 CFU/g), Rushan Visitor Center (up to 516 CFU/g), and Heping Village Park (up to 320 CFU/g). In contrast, agricultural areas and forest sites generally showed lower fungal burdens, typically ranging from 0-200 CFU/g.

Multiple species were often isolated from the same sampling site, particularly in urban areas. Analysis of species co-occurrence revealed that recreational and public areas harbored higher average numbers of coexisting species (0.72 and 0.65 species per site, respectively) compared to agricultural areas (0.32 species per site) and natural forests (0.25 species per site). The most diverse communities were found at Chihshang No3 Park (4 species: *S. apiospermum, S. hainanense, S. minutisporum*, and *S. dehoogii*), followed by Hsinchu Park (3 species) and Taipei Veterans General Hospital Fenglin Branch (3 species). These findings suggest that recreational and public environments might provide more suitable conditions for diverse *Scedosporium* and *Lomentospora* communities compared to agricultural lands and natural forests.

### Fungal Community Dispersion and Land Use Correlations

Analysis of multivariate dispersion revealed differences in fungal community heterogeneity across land usage types. Public and recreational areas showed greater community dispersion (distances to median of 0.607 and 0.560, respectively) compared to agricultural and forest areas (0.541 and 0.537 respectively). However, ANOVA results on the homogeneity of multivariate dispersions confirmed no significant differences among land usage types (F = 0.540, df = 3, 204, p = 0.655), indicating the degree of variation within each land usage type was similar. Tukey’s post-hoc pairwise comparisons showed no significant differences in community dispersion across land usage types (all p-adj > 0.05), with the largest difference between public and forest areas (diff = 0.069, p-adj = 0.471), though still not significant (Figure 2).

**Figure 2.**
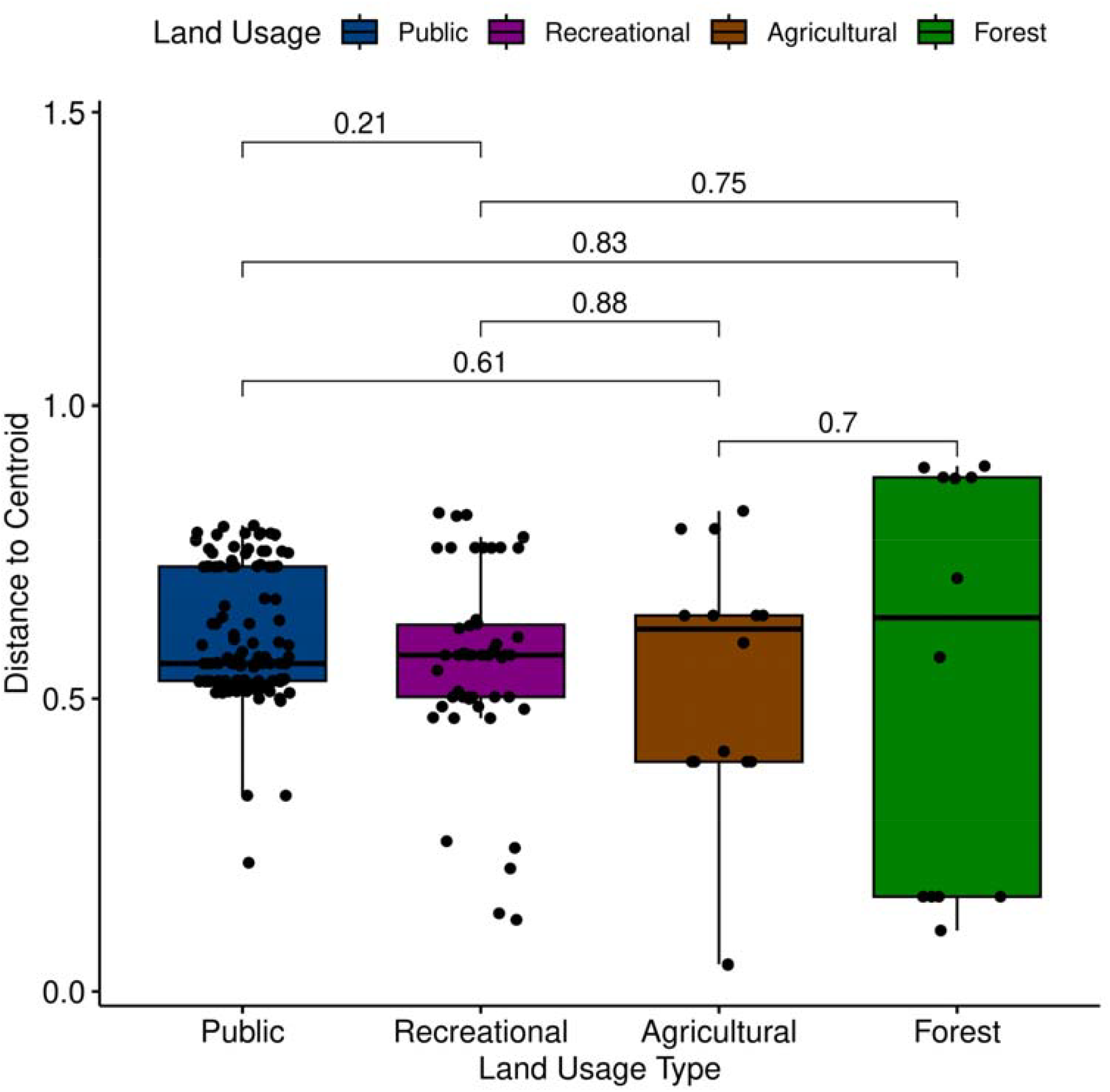
Homogeneity test of species assemblages across different land usage types (Public, Recreational, Agricultural, Forest). The distances to the centroid for each land usage type are shown along the y-axis, with p-values from pairwise comparisons displayed above the bars.

Analysis of soil samples revealed wide variation in chemical parameters across sampling sites. Total nitrogen content ranged from 0.038 to 0.613%, carbon content from 0.398 to 11.363%, and organic content from 0.644 to 19.211%. Water content varied between 1.064 and 10.609%, while pH values ranged from 4.665 to 7.735. Electrical conductivity (EC) fluctuated between 0.360 and 1.215 mS/cm, and available phosphorus showed substantial variation from 2.477 to 324.624 ppm. Base cations exhibited the following ranges: sodium (0.105-0.675%), potassium (0.006-0.157%), magnesium (0.296-4.907%), and calcium (8.110-63.475%). The Human Footprint Index (HFI) of sampling sites ranged from 8 to 71, indicating a broad spectrum of human impact.

Statistical analysis revealed a significant positive correlation between fungal abundance (Mean CFU) and HFI (r = 0.125, p = 0.013), suggesting that human activity influences fungal distribution patterns (Figure 3). Among soil chemical parameters, electrical conductivity showed a weak negative correlation (r = -0.336, p = 0.056), though not statistically significant. Other edaphic factors including total nitrogen (r = -0.119, p = 0.508), carbon content (r = -0.124, p = 0.491), organic content (r = -0.201, p = 0.346), water content (r = 0.006, p = 0.972), pH (r = 0.004, p = 0.983), available phosphorus (r = -0.036, p = 0.840), sodium (r = -0.198, p = 0.268), magnesium (r = -0.246, p = 0.168), calcium (r = -0.016, p = 0.928), and potassium (r = 0.009, p = 0.959) showed minimal correlations with fungal abundance. These results indicate that among the measured parameters, human activity had the strongest influence on fungal distribution patterns while individual soil chemical properties, despite their wide ranges, had limited direct effects on fungal abundance.

**Figure 3.**
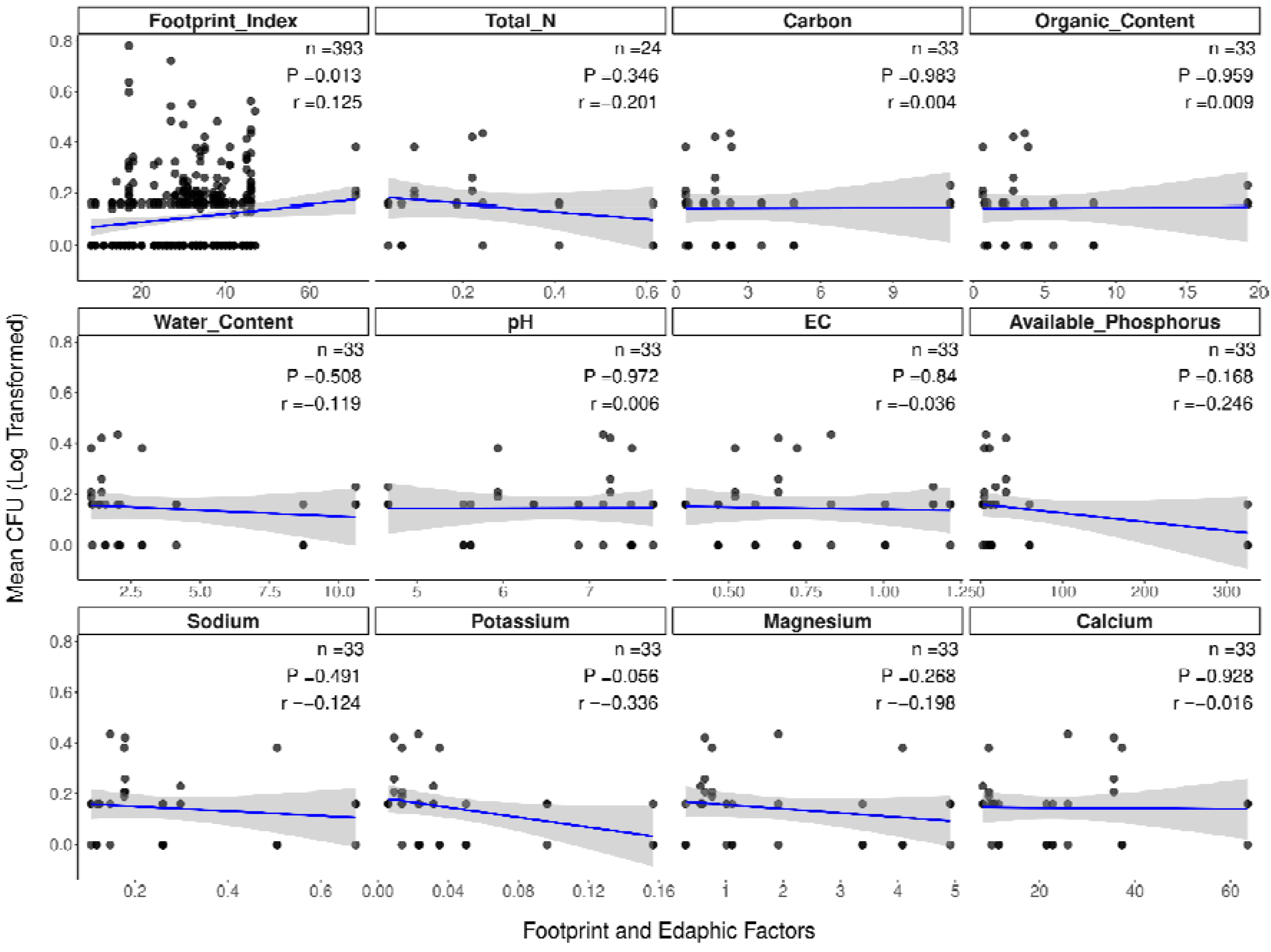
Generalized Linear Model analysis illustrates the relationship between Mean CFU (log-transformed fungal burden) and the Human Footprint Index (HFI) alongside various edaphic chemical factors. Each panel includes the GLM trend line, with annotations for sample size (n), p-value, and correlation (r) for each factor. HFI is the only statistically significant factor influencing fungal abundance, with no significant associations observed for the other soil parameters.

## Discussion

This study offers a comprehensive ecological analysis of *Scedosporium* and *Lomentospora* species distribution across Taiwan, emphasizing the influence of human activity on their habitat preferences. By examining the Human Footprint Index (HFI), soil chemical parameters, and mapping species distributions from 406 diverse soil samples, we identified a strong association between these fungi and anthropogenically disturbed environments.

Contrary to prior assumptions that soil chemistry is the one of the major determinants, our findings highlight human activity as the primary driver of their distribution. This suggests that urbanization and human presence create unique ecological niches conducive to the persistence and proliferation of these species.

### Anthropogenic Environments as Selective Niches for *Scedosporium*/*Lomentospora* Species

Our ecological survey reveals an intriguing pattern of habitat specialization among *Scedosporium* and *Lomentospora* species, with clear preference for anthropogenically modified environments. The highest fungal burdens were detected in urban centers, particularly in intensively used public spaces (Tainan High Speed Rail Station: 580 CFU/g) and healthcare facilities (Taipei Veterans General Hospital: 1293 CFU/g). This distribution pattern suggests these fungi have successfully adapted to exploit niches created by human disturbance of the environment. Global ecological studies consistently demonstrate this anthropogenic habitat preference □ —from European surveys showing these fungi thriving in industrial zones and wastewater treatment plants while being absent from natural forests,^9,10^ to Australian studies documenting higher populations in urban cores versus peripheral areas,^36^ to African surveys finding them concentrated in polluted urban soils,^14^ to Thailand studies revealing their prevalence in high human-traffic areas.^11,12^ This consistency across diverse geographical regions and urban contexts provides strong evidence that these fungi have evolved traits allowing them to exploit conditions characteristic of human-modified environments. The particularly high fungal burdens we observed (up to 1293 CFU/g) exceed previously reported densities, indicating these organisms may find optimal growth conditions in Taiwan’s intensively developed urban environments.

### Biogeographical Patterns Reveal Regional Differences in Species Community

Our survey documents clear geographical variation in *Scedosporium* species composition and relative abundances. In Taiwan’s urban environments, we found *S. boydii* (32.2%) and *S. apiospermum* (30.9%) as the predominant species, followed by *S. dehoogii* (14.4%) and *S. aurantiacum* (8%). This community structure reflects a distinct regional pattern that contrasts with global distributions. European studies reveal varying dominance patterns - French surveys found *S. dehoogii* predominating,^10^ while Austrian investigations showed *S. apiospermum* (69.8%) prevalence.^9^ In the Asia-Pacific region, Thailand reported overwhelming *S. apiospermum* dominance (86.6%),^11^ while Australian studies revealed a unique pattern with *S. aurantiacum* (54.6%) and *L. prolificans* (43%) as the main species.^36^ North African surveys from Morocco demonstrated yet another pattern with *S. apiospermum* dominance but different co-occurring species.^14^ Notably, our study found a more even distribution of species contrary to the single-species dominance reported elsewhere. These geographical variations likely reflect complex interplays between anthropogenic activities (such as urbanization patterns and pollution types), historical biogeographical factors, and perhaps local environmental conditions (including temperature, pH, and nutrient availability).

### Culture-based Detection Reveals Complex Community Structure

Our culture-based approach using DG18 agar achieved a 51.2% positive detection rate, comparable to previous studies using various selective media.^11,18,36^ However, beyond simple detection, our methodology revealed complex patterns of species co-occurrence that provide insights into community assembly. The discovery of up to four coexisting species at single sites (e.g., Chihshang No3 Park harboring *S. apiospermum, S. hainanense, S. minutisporum*, and *S. dehoogii*) suggests these fungi might have evolved mechanisms for resource partitioning that enable coexistence. This level of species co-occurrence exceeds that typically reported in previous studies, which often found only one or two species per site.^10,11,14,36^ While different selective media have been employed across studies (SceSel+, Scedo-Select III), the consistent range of positive culture rates (40-60%) suggests these methods effectively capture the culturable portion of these fungal communities. The higher species co-occurrence we detected may reflect either more sensitive detection methods or genuinely more complex community structures in Taiwan’s urban environments.

### Edaphic Factors Show Limited Influence on Distribution

Our extensive analysis of soil chemical properties revealed complex patterns in their relationship with fungal distribution. Despite examining a wide range of parameters - including total nitrogen (0.038-0.613%), carbon (0.398-11.363%), organic content (0.644-19.211%), and various base cations—we found no significant correlations between these edaphic factors and fungal abundance (Figure 3). Only the HFI showed a significant positive correlation (r = 0.125, p = 0.013), highlighting the predominant influence of anthropogenic activities on fungal distribution patterns.

These findings partially contrast with previous studies. Kaltseis et al. (2009)^9^ reported a positive correlation between ammonium concentration and *Scedosporium* abundance in Austria, with fungi predominantly found in soils with pH 6.1-7.5. Similarly, Luplertlop et al. (2016)^11^ detected these fungi primarily within this pH range in Bangkok. While our study found *Scedosporium*/*Lomentospora* species across a broader pH range (4.665-7.735), aligning with findings in Western France (pH 4.9-8.5),^10^ pH showed no significant correlation with fungal abundance (r = 0.004, p = 0.983). The weak negative trend with potassium (r = - 0.336, p = 0.056) suggests a possible influence of this nutrient on fungal growth, though this relationship requires further investigation.

The lack of strong correlations between soil chemical parameters and fungal distribution, coupled with the significant relationship with human activity, suggests that anthropogenic disturbance of environments is the primary driver of *Scedosporium*/*Lomentospora* distribution. This finding has important implications for understanding infection risks, particularly in urban and industrial areas. The ability of these fungi to thrive across diverse chemical conditions likely reflects their metabolic versatility, including their documented capacity to degrade various organic pollutants.^17,37^

### Human Activity as Primary Driver of Distribution: Quantitative Evidence

Integrating the HFI with traditional analytical methods provides quantitative evidence linking human activity to the distribution of *Scedosporium* and *Lomentospora*. The HFI is constructed from multiple indicators of human pressure—such as built environments, population density, nighttime lights, roads, and navigable waterways—summed to create a composite measure of human influence at a 1 km^2^ resolution.^34^ Our analysis shows a significant positive correlation (r=0.149, p=0.003) between HFI and these fungi, but the modest correlation coefficient suggests human activity explains only a limited portion of the variability in fungal distribution. This advances past studies, which relied primarily on categorical land use [e.g. Kaltseis et al.,(2009);^9^ Harun et al., (2010)^36^], by offering a quantitative measure of human influence. Patterns in species co-occurrence support the influence of human-disturbed environments: recreational and public areas exhibit higher fungal species richness (0.72 and 0.65 species per site) compared to agricultural (0.32) and forested areas (0.25). The relatively low correlation with HFI and lack of associations with measured soil parameters suggest that fungal distribution in these areas may be shaped by a complex mix of factors beyond broad human influence, including community heterogeneity in public spaces. These results underscore that while human activities impact *Scedosporium* and *Lomentospora* distributions, our current approach only partially reveals the environmental dynamics involved.

## Conclusion

Our findings emphasize that human-disturbed environments significantly shape the distribution and diversity of *Scedosporium* and *Lomentospora* species across Taiwan. While the HFI provided a quantifiable link between human activity and fungal distribution, the modest correlation coefficient suggests a need for more specific and relevant indicators to fully capture these patterns. Targeted indicators—such as urban heat intensity, air quality levels, waste density—could offer clearer insights into how anthropogenic factors shape fungal communities. Expanding research into their genomic adaptations and resilience mechanisms could further enhance our understanding of their pathogenicity and inform strategies to mitigate their impact, particularly in healthcare and densely populated urban settings. Together, these refined approaches may improve predictive models and guide more effective public health strategies to manage the risks posed by these emerging pathogens.

